# IL-33-binding HpARI family homologues with divergent effects in suppressing or enhancing Type 2 immune responses

**DOI:** 10.1101/2023.10.09.561567

**Authors:** Florent Colomb, Adefunke Ogunkanbi, Abhishek Jamwal, Beverley Dong, Rick M. Maizels, Constance A. M. Finney, James D. Wasmuth, Matthew Higgins, Henry J. McSorley

**Author notes:** Contributed equally (Florent Colomb, Adefunke Ogunkanbi, Abhishek Jamwal).

## Abstract

HpARI is an immunomodulatory protein secreted by the intestinal nematode *Heligmosomoides polygyrus bakeri*, which binds and blocks IL-33. Here, we find that the *H. polygyrus bakeri* genome contains 3 HpARI family members, and that these have different effects on IL-33-dependent responses in vitro and in vivo, with HpARI1+2 suppressing, and HpARI3 amplifying these responses. All HpARIs have sub-nanomolar affinity for mouse IL-33, however HpARI3 does not block IL-33-ST2 interactions. Instead, HpARI3 stabilises IL-33, increasing the half-life of the cytokine and amplifying responses to it in vivo. Together these data show that *H. polygyrus bakeri* secretes a family of HpARI proteins with both overlapping and distinct functions, comprising a complex immunomodulatory arsenal of host-targeted proteins.

## Introduction

Parasitic helminth infection affects more than one billion people worldwide. The relationship between parasites and their hosts reflects co-evolutionary adaptation to host protective immune mechanisms which results in parasites’ long persistence in the host (1, 2). Excretory-secretory (ES) products released from helminths contain a wide range of molecules which can modulate the host immune system, suppressing anti-parasitic immune responses and hence promote parasite survival (3, 4). The murine intestinal parasite *Heligmosomoides polygyrus bakeri* (*Hpb*) is a prime example of this host-parasite dynamic. *Hpb* secretions contain the Alarmin Release Inhibitor (HpARI) and Binds Alarmin Receptor and Inhibits (HpBARI), both of which block the IL-33 pathway and suppress type 2 immune responses in mice (5, 6). *Hpb* infects mice when larvae are ingested. These larvae rapidly penetrate the epithelium of the duodenum, and develop to adults which emerge into the gut lumen at around day 10 of infection, living for weeks as sexually mature adults in the small intestine. The early phases of infection are associated with damage to the intestinal epithelium as it is penetrated by migrating infective larvae (7).

IL-33 is an IL-1 family alarmin cytokine which is constitutively expressed by epithelial cells, where it is stored in cellular nuclei and released under conditions of damage and necrosis (8). Upon release, the active mature cytokine is rapidly oxidised, rendering it unable to bind to its receptor ST2 (9), limiting its ST2-dependent activity in vivo to the local milieu and to a short period of time after release. IL-33 acts as an initiator of allergic diseases, especially asthma where both IL-33 and ST2 are genetically linked to asthma heritability (10), and are targets for new biologic treatments (11, 12). IL-33 also plays a significant role in parasite expulsion, with deficiency or blockade of the IL-33 pathway resulting in increased parasite burden (8, 13, 14). In our previous work we identified HpARI, showing that it binds IL-33 and blocks type 2 immune responses (5), and used structural studies to demonstrate how this IL-33-ST2 inhibition is achieved (15). Here, we show that the *Hpb* genome encodes a family of three HpARI proteins: HpARI1, HpARI2 and HpARI3, with the original HpARI now renamed as HpARI2. We expressed and tested these proteins for their activity in vitro and in vivo, and unexpectedly found that HpARI3 enhances rather than suppresses IL-33 responses.

## Methods

### Mass spectrometry

The excretory/secretory products of *Hpb* (HES) was prepared as described previously (16), and a 2 μg sample was used for mass spectrometry analysis. Triethylammonium bicarbonate (Sigma) and dithiothreitol (Merck Millipore) were added to final concentrations of 100 mM and 10 mM, respectively, and sample incubated at 70°C for 10 min. Alkylation was carried out by addition of 20 mM iodoacetamide for 30 min in the dark. Sample was incubated overnight in the presence of 50 mM dithiothreitol and 200 ng trypsin was added, and incubated overnight at 37°C. Trifluoracetic acid (Fisher Scientific) at 10% sample volume was added, C18 clean-up was carried out using an Empore-C18 (Agilent) solid phase extraction cartridge, and dried samples resuspended in 1% Formic acid (Fisher Chemical).

Analysis of peptide readout was performed on a Q Exactive™ plus Mass Spectrometer (Thermo Scientific) coupled to a Dionex Ultimate 3000 RS (Thermo Scientific). Liquid chromatography buffers used were: buffer A (0.1% formic acid in Milli-Q water (v/v)) and buffer B (80% acetonitrile and 0.1% formic acid in Milli-Q water (v/v). An equivalent of 1.0 μg of peptides from each sample were loaded at 10 μL/min onto a μPAC trapping C18 column (Pharmafluidics). The trapping column was washed for 6 min at the same flow rate with 0.1% TFA and then switched in-line with a Pharma Fluidics, 200 cm, μPAC nanoLC C18 column. The column was equilibrated at a flow rate of 300 nl/min for 30 min. The peptides were eluted from the column at a constant flow rate of 300 nl/min with a linear gradient from 2% buffer B to 5.0% buffer B in 5 min, from 5.0% B to 35% buffer B in 125 min, and from 35% to 98% in 2 minutes. The column was then washed at 98% buffer B for 20 min and then washed at 2% buffer B for 20 minutes. Two blanks were run between each sample to reduce carry-over. The column was kept at a constant temperature of 50°C.

Q-exactive plus was operated positive ionization mode using an easy spray source. The source voltage was set to 2.90 kV and the capillary temperature was 275°C. Data were acquired in Data Independent Acquisition Mode as previously described (17), with little modification. A scan cycle comprised a full MS scan (m/z range from 345-1155), resolution was set to 70,000, AGC target 1x10^6^, maximum injection time 100 ms. MS survey scans were followed by DIA scans of dynamic window widths with an overlap of 0.5 Th. DIA spectra were recorded at a resolution of 17,500 at 200 m/z using an automatic gain control target of 2x10^5^, a maximum injection time of 100 ms and a first fixed mass of 100 m/z. Normalised collision energy was set to 27% with a default charge state set at 3. Data for both MS scan and MS/MS Data Independent Acquisition (DIA) scan events were acquired in profile mode.

Mass spectrometry data analysis: Label-free analysis was performed in Maxquant (version 2.0.3.0) using the generate RAW files. Enzyme specificity was set to that of trypsin, allowing for cleavage N-terminal to proline residues and between aspartic acid and proline residues. Other parameters used were: (i) variable modifications-oxidation (M), protein N-acetylation, gln → pyro-glu, deamidation (NQ), deoxidation (MW); (ii) fixed modifications, cysteine carbamidomethylation; (iii) database: in-house database; (iv) MS/MS tolerance: FT-MS 10ppm, FT-MSMS 0.06 Da; (v) maximum peptide length, 6; (vi) maximum missed cleavages, 2; (vii) maximum of labelled amino acids, 3; and (vii) false discovery rate, 1%. Match Between Runs (MBR) was set to true. Unique peptides were used for protein quantification.

### Expression and Purification of HpARI1-3

Proteins were expressed and purified using a protocol described previously (5). Briefly, mammalian expression constructs carrying C-terminal 6X-His tagged gene of interest (HpARI1-3, or ST2 ectodomain) were transfected individually into Expi293F cells (Thermo Fisher) using the Expifectamine transfection kit (Thermo Fisher). Cell supernatants were harvested 96 hours post-transfection and the expressed recombinant proteins were than captured from the filtered supernatants using Ni-NTA chromatography.

### Surface Plasmon Resonance

To measure their binding affinities for mIL-33, the purified HpARI variants were first polished to remove aggregates, using a Superdex S75 10/300 column in 1XPBS and chemically biotinylated using EZ-Link™ Sulfo-NHS-Biotin (Thermo Fisher Scientific) following manufacturer’s instruction. All SPR experiments were carried out using a Biacore T200 instrument (GE Healthcare) using a CAP chip (Biotin CAPture kit (Cytiva)) in an SPR buffer containing 20 mM Tris-Cl pH 8.0, 200 mM NaCl, 1mg/ml salmon sperm DNA and 0.05% Tween-20. Purified mIL-33 was equilibrated in the SPR buffer using a PD-5 column prior to the experiment. The sensor surface was first coated with oligonucleotide coupled streptavidin following manufacturer’s instructions. Individual biotinylated HpARI variants were captured on different flow cells followed by individual injections of 2-fold concentration series of mIL-33 (50 nM to 0.195 nM for HpARI1 and HpARI2 and 200 nM to 0.781 nM for HpARI3) at a flow rate of 40 μl/min, with 400 s association time and 1000 s dissociation time. After each injection, the sensor chip surface was regenerated by injecting 10 μl of 6M guanidium-HCl and 1M NaOH pH 11.0 mixed in a 4:1 ratio. All SPR data were analysed using the BIA evaluation sonware 2.0.3 (GE Healthcare).

### Animals

BALB/cAnNCrl and C57BL/6JCrl mice were purchased from Charles River, UK. Heterozygous IL-13eGFP mice were provided by Prof Andrew McKenzie (18) and were bred in-house. Experiments were caged-blocked: each cage contained one member of each group in the experiment, thus controlling for cage effects. Mouse accommodation and procedures were performed under UK Home Office licenses with institutional oversight performed by qualified veterinarians.

### In vivo Alternaria model challenge

BALB/c mice were intranasally administered with 50 μg Alternaria allergen (Greer XPM1D3A25) and 10 μg HpARI1, HpARI2 or HpARI3 suspended in PBS, carried out under isoflurane anaesthesia. Mice were culled 24 h later and bronchoalveolar lavage (BAL) was collected (4 lavages with 0.5 ml ice-cold PBS). Lungs were taken for single-cell preparation and flow cytometry, as previously described (5). Lung and BAL eosinophils were identified as SiglecF^hi^CD11c^−^CD45^+^ live cells. Lung ILC2 were identified as ICOS^+^CD90.2^+^Lineage^−^CD45^+^ cells. IL-5 levels were quantified in undiluted BAL fluid by ELISA following manufacturer’s instructions (Invitrogen).

### CMT-64 culture and treatment

CMT-64 cells (ECACC 10032301) were maintained in complete RPMI [RPMI 1640 medium containing 10% fetal bovine serum, 2 mM L-glutamine, 100 U/ml Penicillin and 100 μg/ml Streptomycin (ThermoFisher Scientific)] at 37°C, 5% CO_2_. 96-well plates were seeded at 5x10^4^ cells per well. Cells were grown to 100% confluency, washed and incubated in complete RPMI containing HpARI2 or HpARI3 at 10 μg/ml or in combination at various concentration as indicated. Cells were snap frozen on dry ice for at least 1 h, then thawed and incubated at 37°C as indicated, prior to collection of supernatants and application to bone marrow cell cultures.

### Bone marrow assay and ELISA

Bone marrow single cell suspensions were prepared from C57BL/6 mice, by flushing tibias and femurs with RPMI 1640 medium using a 21 g needle. Cells were resuspended in ACK lysis buffer (Gibco) for 5 min at room temperature, prior to resuspension in complete RPMI (with 10% FCS, 1% Penicillin/Streptomycin, 1% L-glutamine, Gibco) and passing through a 70 μm cell strainer. Cells were cultured in round-bottom 96-well-plates in a final 200 μl volume, containing 0.5×10^6^ cells/well. IL-2 and IL-7 (Biolegend) were added at 10 ng/ml final concentration. CMT freeze-thaw supernatant prepared as described above were added at 50 μL per well. Cells were then cultured at 37°C, 5% CO2, for 5 days, prior to supernatant collection. IL-5 and IL-13 concentration were assessed following manufacturer’s instructions using mouse uncoated IL-5 and IL-13 ELISA kits (Invitrogen).

### Assessment of ST2 binding by IL-33 in the presence of HpARI variants

Purified mST2 ectodomain (20 μM) was mixed individually with 40 μM of mIL33, 25 μM HpARI2:mIL33 and HpARI3:mIL33 to a final volume of 200 μl. All samples were incubated for 1 hour on ice and then applied to the Superdex S200 10/300 GL column (cytiva) pre-equilibrated with buffer containing 20 mM Tris-Cl pH 8.0 and 100 mM NaCl. The eluted fractions were analyzed by SDS-PAGE.

### Statistics

Data was analysed using Graphpad Prism v10.0.0. When comparing independent groups, one-way ANOVA with Dunnet’s post test was used. When comparing groups over a timecourse, two-way ANOVA with Dunnet’s post test was used. Standard error of mean is used throughout. **** = p < 0.0001, *** = p < 0.001, ** = p < 0.01, * = p < 0.05, N.S. = Not significant (p > 0.05).

## Results

### *H. polygyrus bakeri* genome encodes three HpARI-like proteins

HpARI was identified by in-house transcriptomic and proteomic analysis of *Hpb* (5). Homology searching of published *Hpb* transcriptomic and genomic data via Wormbase ParaSite (19) identified a family of 3 HpARI genes from *Hpb*, all of which have the same characteristic domain organisation of a secretory signal peptide followed by 3 Complement Control Protein (CCP) domains (**Figure 1A**). Proteomic analysis of *Hpb* excretory/secretory products (HES) showed that the most abundant HpARI family member was HPOL_0002204701 (Intensity = 9x10^9^), and so was named HpARI1, while the previously-identified “HpARI” protein, HPBE_0000813301 (5), was renamed as HpARI2 (Intensity = 1x10^9^), and the final homologue with lowest intensity in HES, HPBE_0002531401, was renamed HpARI3 (Intensity = 4x10^7^). Further analysis of a recent transcriptomic study of *Hpb* (20) showed that while HpARI1 and HpARI3 are expressed at a fairly consistent level throughout the lifecycle (albeit with HpARI3 at a far lower level compared to HpARI1), the expression of HpARI2 peaks early in infection (during the tissue-dwelling phase), with similar trends in males and females for each HpARI (**Figure 1B**). The 3 HpARI family members show a high level of sequence similarity (69-81% identity between the 3 HpARIs), however the HpARI3 sequence contains a notable 3 amino acid deletion (at E94_R96 in HpARI2 sequence) at the N-terminal end of CCP2 (**Figure 1A**), a region which contains an HpARI2-IL-33 interaction site (15).

**Fig 1:**
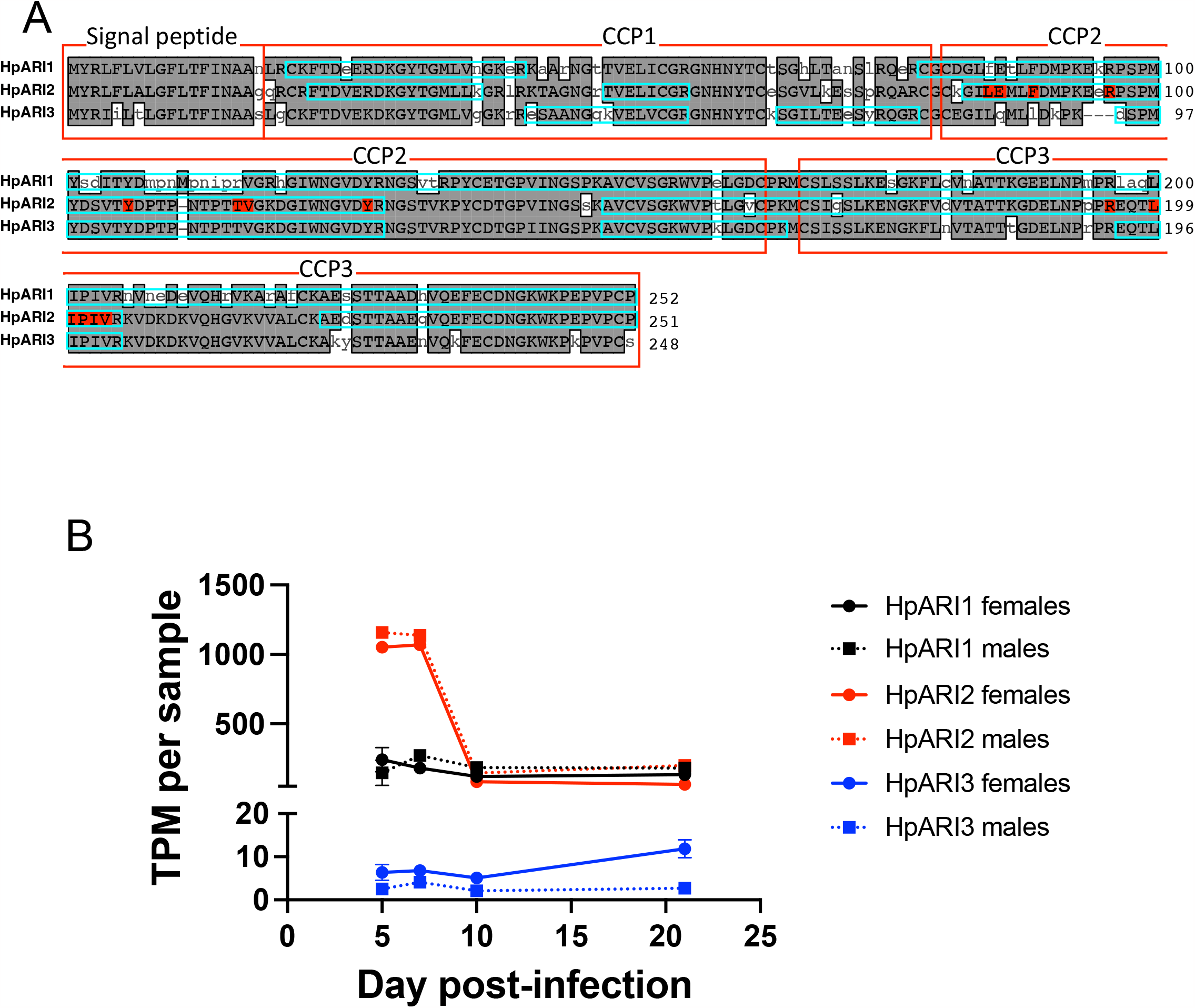
Sequences of HpARI family members. A. Alignment of HpARI1, HpARI2 and HpARI3. Grey highlight indicates agreement with consensus. Signal peptide and CCP domains are indicated by red boxes, coverage of peptides identified by mass spectrometry of HES indicated by blue boxes. Residues in HpARI2-IL-33 interface highlighted in red. B. Transcription of HpARI1, HpARI2 and HpARI3 in male and female parasites over a time course of infection. Data retrieved from the Sequence Read Archive, PRJNA750155.

### The HpARI family members all bind IL-33 with sub-nanomolar affinities

The 3 HpARI family members were expressed in mammalian cells and purified for further testing. Each HpARI was added to the CMT-64 cells (which express high levels of IL-33 (5)), followed by freeze-thaw necrosis-induced IL-33 release. In this assay, all 3 proteins could suppress the IL-33 signal in a dose dependent manner, indicating that they compete for IL-33 binding with the ELISA antibodies used for detection (**Figure 2A**). HpARI3 required an approximately 8-fold higher concentration for IL-33 suppression (IC50 of HpARI1 and HpARI2 ∼8x10^−10^ M, HpARI3 ∼6x10^−9^ M). When binding affinity for IL-33 was tested by surface plasmon resonance, HpARI3 was likewise shown to have a lower affinity for IL-33, albeit still with a sub-nanomolar K_D_: HpARI1/2 have K_D_ around 4x10^−11^ M, while HpARI3 has a K_D_ of 4x10^−10^ M (**Figure 2B**).

**Figure 2:**
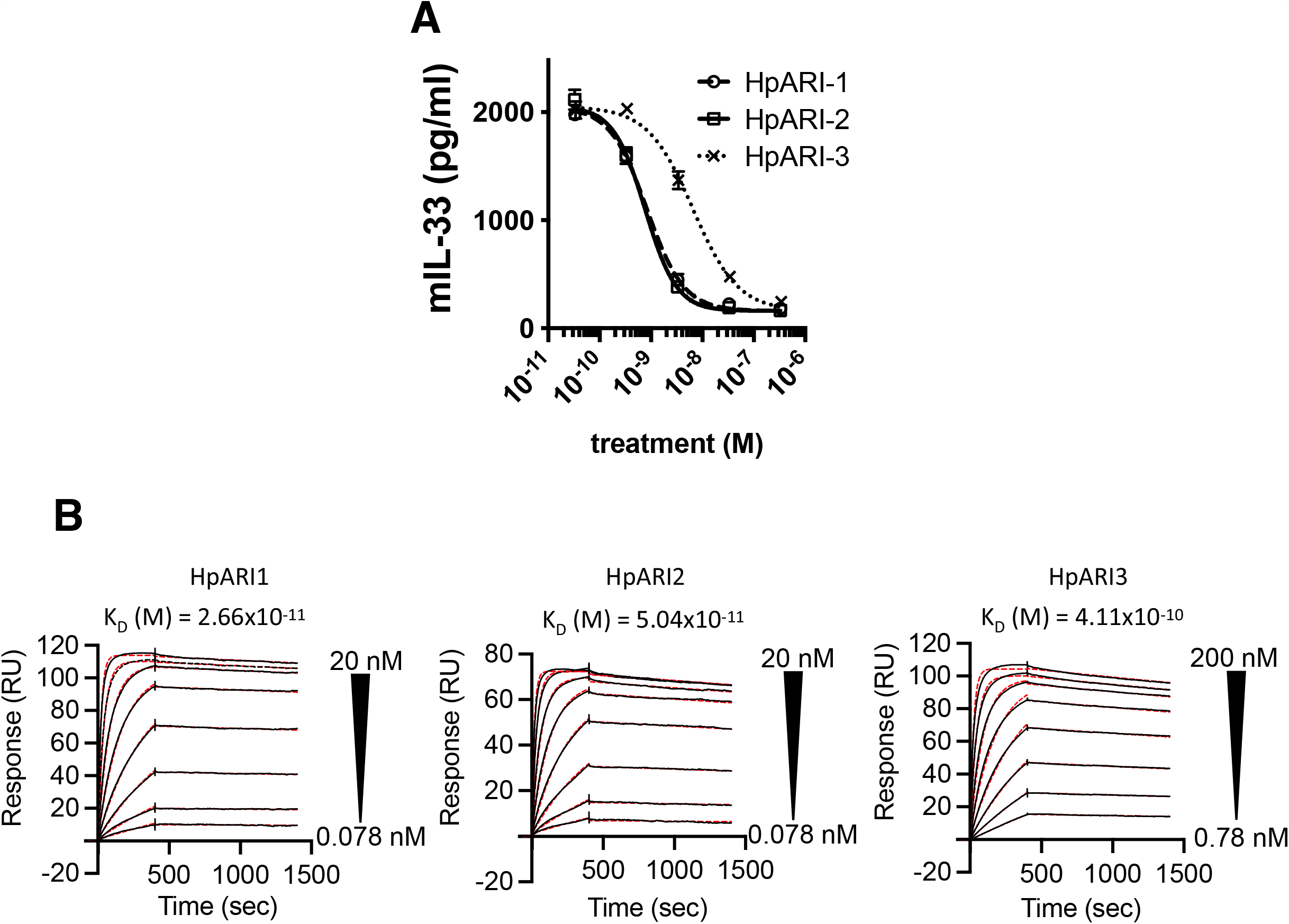
HpARI family members bind IL-33. A. IL-33 detected in supernatant of CMT-64 cultures after freeze-thaw in the presence of HpARI1, HpARI2 or HpARI3. B. Surface plasmon resonance (SPR) analysis of mouse IL-33 binding to chip-bound HpARI2 or HpARI3.

### HpARI family show differing effects on IL-33-dependent responses in vivo

To test how each HpARI affected IL-33-dependent responses, we coadministered them intranasally to BALB/c mice with Alternaria allergen (a highly IL-33-dependent model (21-23)), and assessed type 2 innate lymphoid cell (ILC2) and eosinophil responses 24 h later (**Figure 3A**). HpARI2 potently suppressed BAL and lung eosinophilia, ILC2 responses and IL-5 release, as shown previously (5), while HpARI1 had a similar, albeit slightly blunted, suppressive effect (**Figure 3B-G**). Surprisingly, HpARI3 had the opposite effect, amplifying responses in this model, with significantly increased lung eosinophilia and ILC2 activation (CD25 upregulation and increased cell size measured by FSC) as well as total IL-5 release (**Figure 3B-G**). These increased responses in the presence of HpARI3 are remarkably similar to that seen with a truncation of HpARI2 lacking the CCP3 domain (HpARI_CCP1/2), which also amplifies IL-33 responses in vivo (14, 24). While both HpARI2 full-length protein and the HpARI_CCP1/2 truncation can both bind to IL-33, only HpARI2 full-length protein can block IL-33-ST2 interactions, while HpARI_CCP1/2 cannot (24), therefore we also tested the IL-33 blocking ability of HpARI3.

**Figure 3:**
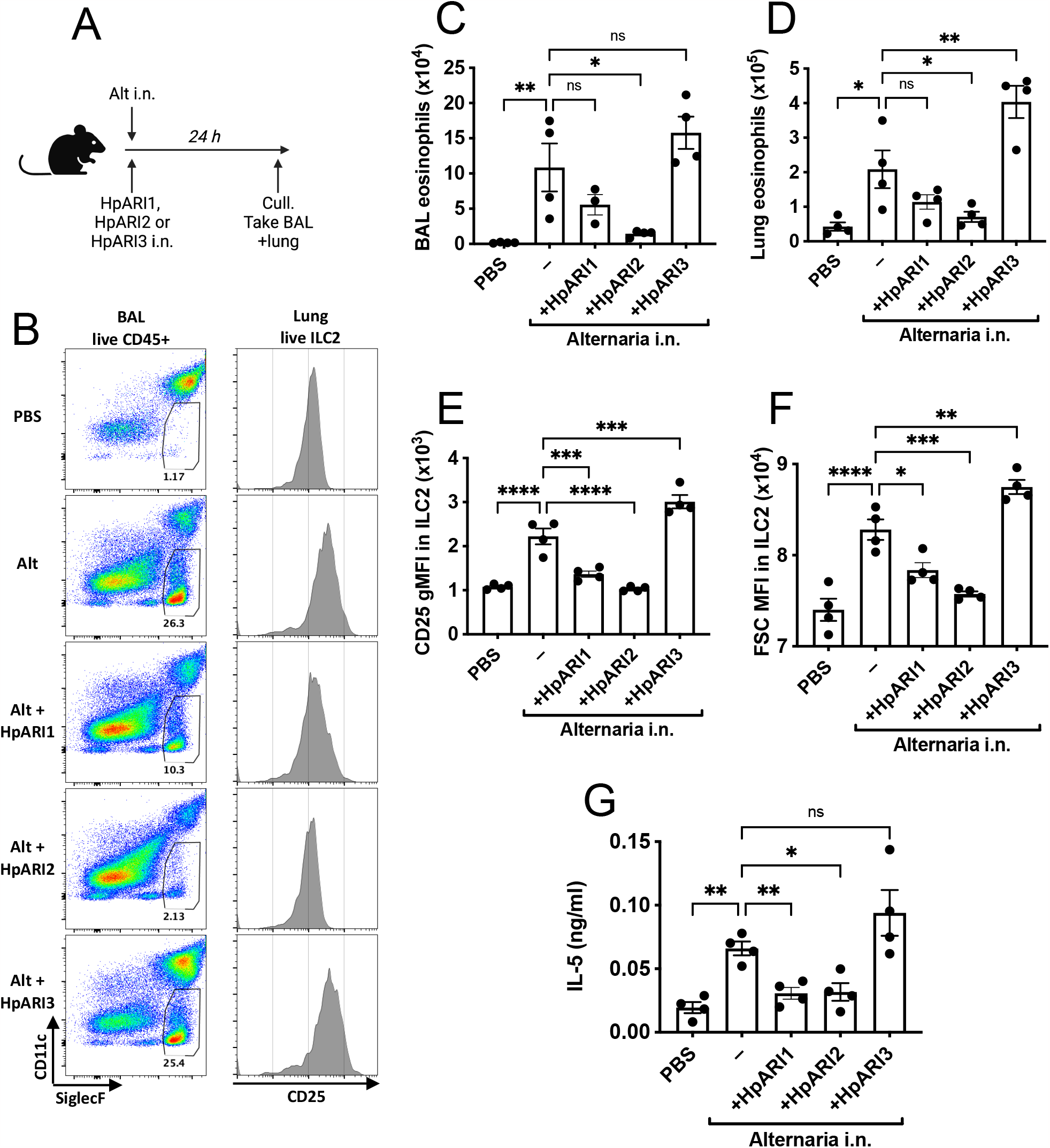
HpARI family members have differing effects against IL-33. A. Experimental setup for B-G. B. Flow cytometry of BAL Siglecf versus CD11c, gated on CD45^+^ live cells, showing eosinophil gate (left). Flow cytometry of CD25 on lung ILC2, gated on live ICOS^+^CD90^+^Lin^−^CD45^+^ cells (right). Representative samples shown. C *Alternaria* allergen was co-administrered with HpARI1, HpARI2 or HpARI3 as shown in A, and BAL eosinophil numbers (Siglecf^+^CD11^−^CD45^+^) measured 24h later D. Eosinophil (Siglecf^hi^CD11^−^CD45^+^) numbers in lung tissue. E. CD25 expression level on lung ILC2 (ICOS^+^CD90^+^Lin^−^CD45^+^). F. FSC mean in lung ILC2 (ICOS^+^CD90^+^Lin^−^CD45^+^) G. IL-5 levels in cell-free BAL fluid. Data representative of two repeat experiments with 4 mice per group. Error bars show SEM.

### HpARI3 does not prevent IL-33 from binding to ST2

To determine whether different HpARI variants can prevent IL-33 from binding to ST2, we deployed a size exclusion chromatography assay to assess IL-33-HpARI-ST2 complex formation (15). We first confirmed the formation of a complex of ST2 and IL-33 as the two proteins co-migrated as a higher molecular weight complex when compared with individual components (**Figure 4A**). In contrast, as previously observed (15) the elution profile of ST2 was unaltered in presence of HpARI2:mIL33 complex, showing that HpARI2 prevents IL-33 from binding to ST2 (**Figure 4B**). In contrast, HpARI3 did not prevent IL-33 from binding to ST2 as indicated by elution peak shift in which ST2, IL-33 and HpARI3 co-elute, showing the formation of a complex containing all three components (**Figure 4C**). Therefore, in contrast to HpARI2, HpARI3 interacts with IL-33 in a manner which does not prevent IL-33-ST2 interaction.

**Figure 4:**
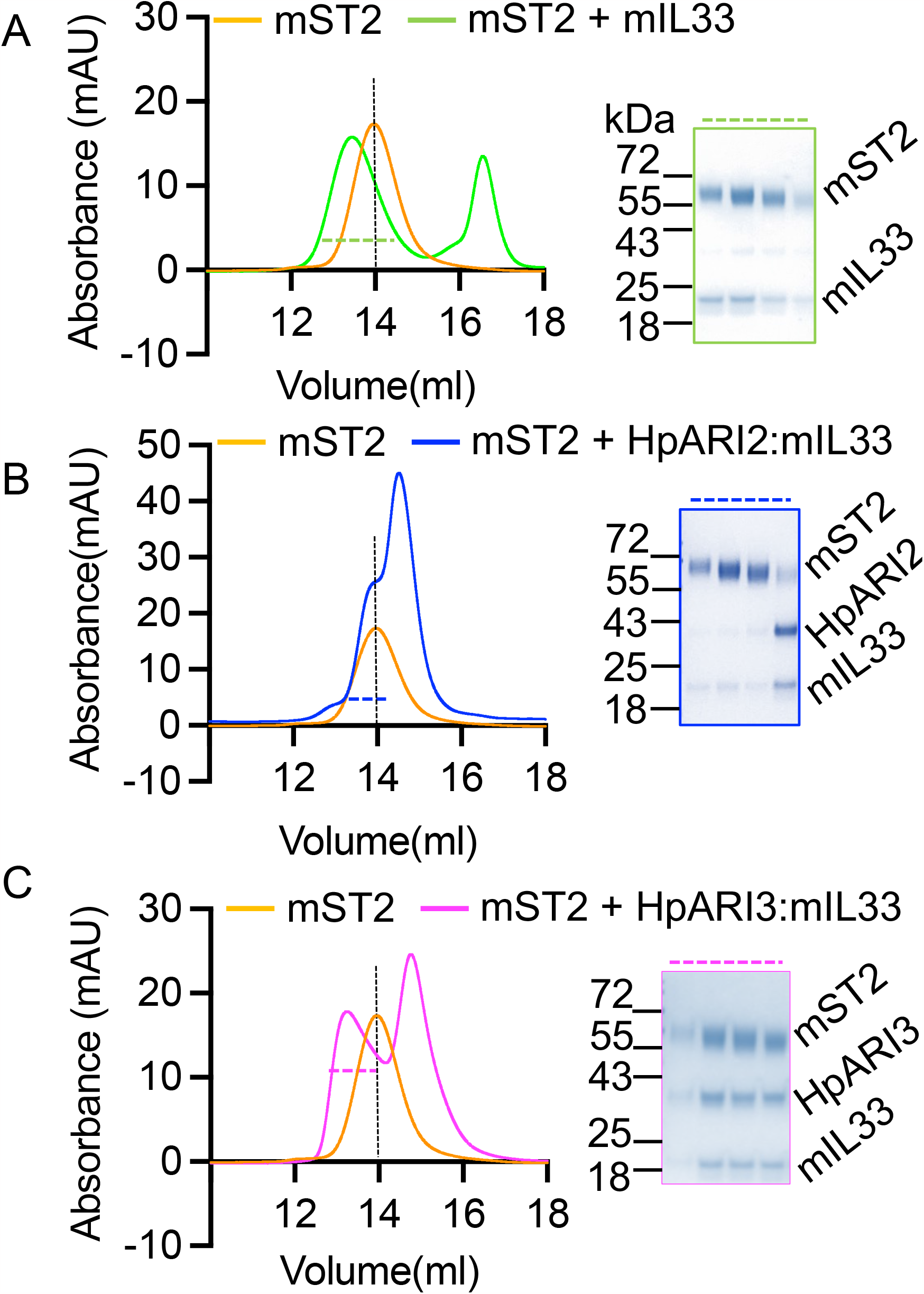
HpARI3 binding does not prevent IL-33-ST2 interaction. A. SEC and SDS PAGE analysis of purified mST2 ectodomain (orange curve) alone, or in presence of mIL33 (green) B. SEC and SDS PAGE analysis of purified mST2 ectodomain (orange curve) alone, or in presence of HpARI2:mIL33 (blue) C. SEC and SDS PAGE analysis of purified mST2 ectodomain (orange curve) alone, or in presence of HpARI3:mIL33 (magenta) SDS-PAGE analysis shows fractions indicated by dotted line in SEC figures.

### HpARI3 stabilises IL-33

As HpARI3 cannot block IL-33-ST2 interactions, and instead amplified responses in an IL-33-dependent model, we hypothesised that HpARI3 binding to IL-33 may stabilise the cytokine, extending its half-life. To test this, we used an in vitro IL-33 release and inactivation assay which we developed previously (24). In this assay, CMT-64 cells are freeze-thawed to induce necrosis, and released IL-33 is incubated at 37°C for between 15 min and 48 h after release, to allow inactivation of the cytokine. Supernatants are subsequently applied to naïve IL-13eGFP murine bone marrow cells in the presence of IL-2 and IL-7 to support ILC2 differentiation, and ILC2 activation was assessed by flow cytometry for activation markers (IL-13eGFP and CD25 upregulation) and ELISA for secretion of IL-5 and IL-13. In medium control wells, CMT-64 freeze-thaw supernatants could only induce significant IL-5 and IL-13 production when applied within 6 h of thaw, indicating inactivation of IL-33 at later points of this timecourse, as shown previously (24). Conversely, CMT-64 cells freeze-thawed in the presence of HpARI2 could not induce any ILC2 activation, due to blockade of released IL-33. In the presence of HpARI3 however, IL-33 in supernatants was able to stimulate high levels of IL-5 and IL-13 release and ILC2 activation (**Figure 5B-E**), indicating strong IL-33 responses which were not inactivated even after incubation of IL-33-containing CMT-64 supernatants for 48 h at 37°C.

**Figure 5:**
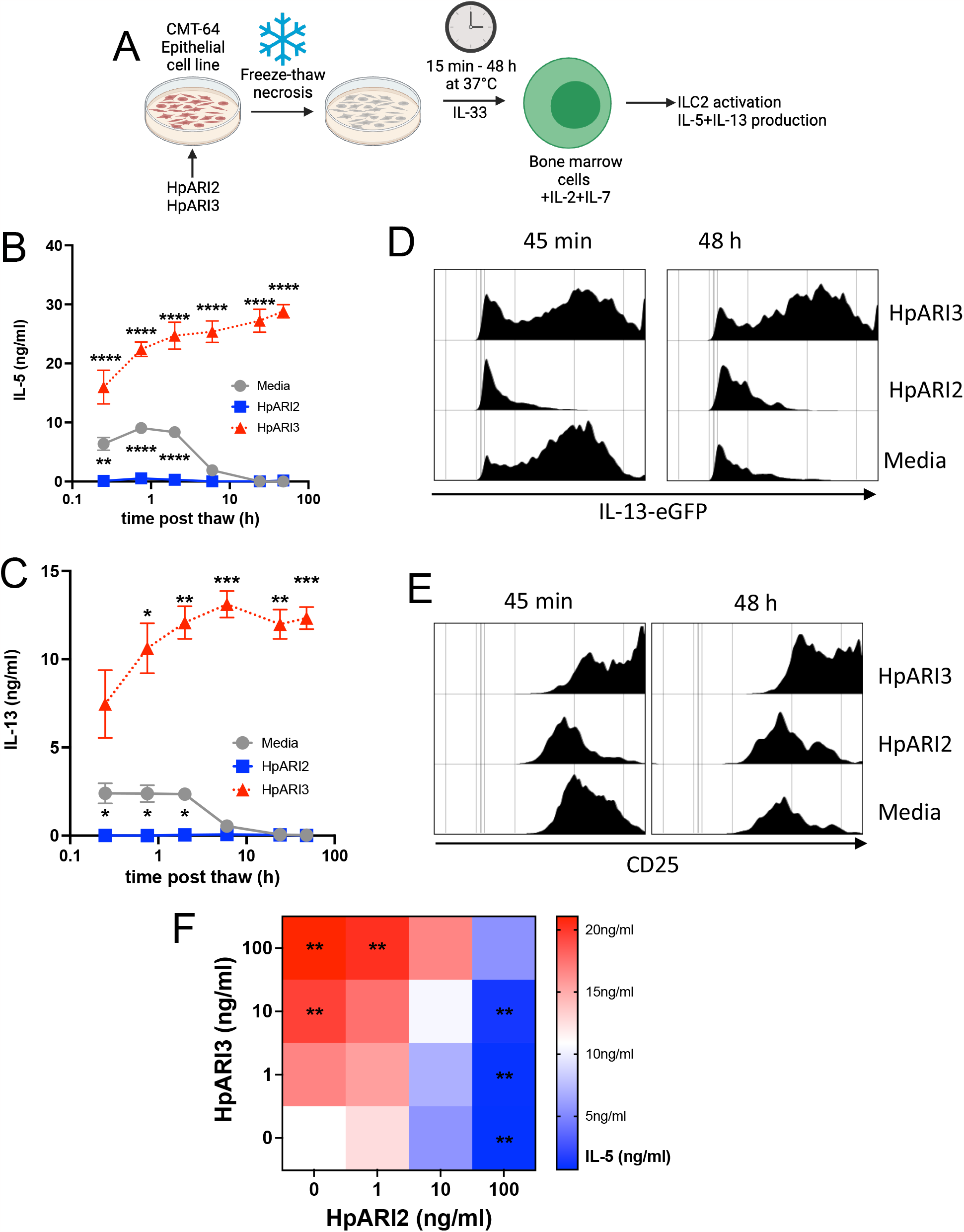
HpARI3 stabilises IL-33 in its active form. A. Experimental setup for B-E. Supernatants were taken over a time course after freeze-thaw of CMT-64 cells in the presence of HpARI2 or HpARI3, transferred to IL-13eGFP^+/-^ bone marrow cells with IL-2+IL-7, and cultured for 5 days. Created with BioRender.com B. IL-5 levels in supernatants from cultures as shown in A, measured by ELISA. C. IL-13 levels in supernatants from cultures as shown in A, measured by ELISA. D. IL-13eGFP expression in gated ICOS^+^Lin^−^CD45^+^ ILC2s from cultures as shown in A, measured by flow cytometry. E. CD25 expression in gated ICOS^+^Lin^−^CD45^+^ ILC2s from cultures as shown in A, measured by flow cytometry. Data from B-E representative of three repeat experiments. Error bar shows SEM of 4 technical replicates. Analysed by 2 way ANOVA with Dunnet’s post test. F. HpARI2 and HpARI3 at a range of concentrations were applied to CMT-64 cells, prior to freeze-thaw, incubation at 37°C for 2 h, and transfer to cultures of bone marrow cells with IL-2+IL-7, and cultured for 5 days. Levels of IL-5 in supernatants were measured by ELISA. Mean of 3 independent biological replicates. Statistical significance shown as compared to medium-only control. Analysed by 1 way ANOVA with Dunnet’s post test.

Finally, to determine whether the effects of HpARI2 and HpARI3 could compete in combination, we added each HpARI at 1, 10 or 100 ng/ml to the assay described above, and measured released IL-5. While sole HpARI3 treatment caused a dose-dependent increase in IL-5 release, HpARI2 caused a decrease, as expected. In combination, equivalent concentrations of each HpARI showed no significant change compared to medium-only control, while an excess of HpARI2 significantly suppressed responses, and an excess of HpARI3 significantly enhanced responses (**Figure 5F**). Therefore, the suppression or enhancement of IL-33 responses in vivo by HpARI2 or HpARI3 will depend on the predominance of each molecule in the local milieu.

## Discussion

In this study, we describe a family of 3 HpARI proteins, with high sequence similarity but differing effects on responses to IL-33. While HpARI1 and HpARI2 suppress responses to IL-33, HpARI3 lacked the ability prevent IL-33 from binding to ST2 and instead stabilised this highly labile cytokine in its active form, resulting in amplification of responses to IL-33 in in vivo and in vitro models.

The IL-33 activity-amplifying ability of HpARI3 was unexpected, and would seem counterintuitive as total ST2-deficient mice are more susceptible to *Hpb* infection (13), implying the IL-33 pathway is largely helminth-toxic. However, additionally to the well-described type 2 immunity-inducing effects of IL-33, recent publications have described a range of effects of IL-33 which can suppress type 2 immunity. IL-33 can activate and expand Foxp3^+^ regulatory T cells (25), and in helminth infection deletion of IL-33 in myeloid cells results in a defective regulatory T cell response, increased type 2 immunity, and accelerated helminth ejection (26). Furthermore, IL-33 can activate natural killer cell (27), Th1 (28) and CD8+ T cell (29) IFN-γ production, potentially suppressing type 2 immunity. As both Th1/CD8+ IFN-γ (30, 31) and Foxp3+ regulatory T cell (32) responses have been demonstrated to control type 2 immunity in *Hpb* infection, either of these pathways could be the target of HpARI3 responses.

Although HpARI3 expression was maintained at a lower level than that of HpARI1 or HpARI2, all 3 HpARI proteins were detectable in HES. As HpARI2 and HpARI3 can compete in in vitro cultures, we hypothesise that differential timing or localisation of HpARI protein release could determine the local HpARI2 / HpARI3 ratio, and therefore the resulting response to IL-33 during infection. IL-33’s inflammatory versus immunosuppressive effects are partially controlled by context of release: when IL-33 was deleted in epithelial cells, type 2 immunity and parasite ejection was deficient, while when IL-33 was deleted in myeloid cells, regulatory T cell responses were reduced, resulting in increased type 2 immunity and acerated parasite ejection (26). It would therefore be most advantageous for the parasite to block IL-33 in the local environment of the parasite during the early tissue-dwelling phase of infection (where necrotic damage to the epithelium is most likely), while IL-33 enhancement in the draining lymph node could amplify Th2-suppressive IFN-γ and regulatory T cell responses. Further work is required to investigate the roles of this family of immunomodulatory proteins during active infection.

## Acknowledgements

This work was funded by awards to HJM from LONGFONDS|Accelerate as part of the AWWA project, the Medical Research Council (MR/S000593/1) and Wellcome 221914/Z/20/Z. We would like to acknowledge the support of Cara Henderson and Alan Score of the FingerPrints Proteomics Facility at the University of Dundee, which is supported by the ‘Wellcome Trust Technology Platform’ award [097945/B/11/Z].

## References

1. Brooker S, Clements AC, Bundy DA. 2006. Global epidemiology, ecology and control of soil-transmitted helminth infections. Advances in parasitology 62:221–261.

2. Hotez PJ, Brindley PJ, Bethony JM, King CH, Pearce EJ, Jacobson J. 2008. Helminth infections: the great neglected tropical diseases. The Journal of clinical investigation 118:1311–1321.

3. Lothstein KE, Gause WC. 2021. Mining Helminths for Novel Therapeutics. Trends in Molecular Medicine 27:345–364.

4. Maizels RM, McSorley HJ. 2016. Regulation of the host immune system by helminth parasites. Journal of Allergy and Clinical Immunology 138:666–675.

5. Osbourn M, Soares DC, Vacca F, Cohen ES, Scott IC, Gregory WF, Smyth DJ, Toivakka M, Kemter AM, le Bihan T, Wear M, Hoving D, Filbey KJ, Hewitson JP, Henderson H, Gonzalez-Ciscar A, Errington C, Vermeren S, Astier AL, Wallace WA, Schwarze J, Ivens AC, Maizels RM, McSorley HJ. 2017. HpARI Protein Secreted by a Helminth Parasite Suppresses Interleukin-33. Immunity 47:739–751 e5.

6. Vacca F, Chauche C, Jamwal A, Hinchy EC, Heieis G, Webster H, Ogunkanbi A, Sekne Z, Gregory WF, Wear M, Perona-Wright G, Higgins MK, Nys JA, Cohen ES, McSorley HJ. 2020. A helminth-derived suppressor of ST2 blocks allergic responses. Elife 9.

7. Gentile ME, Li Y, Robertson A, Shah K, Fontes G, Kaufmann E, Polese B, Khan N, Parisien M, Munter HM, Mandl JN, Diatchenko L, Divangahi M, King IL. 2020. NK cell recruitment limits tissue damage during an enteric helminth infection. Mucosal Immunol 13:357–370.

8. McSorley HJ, Smyth DJ. 2021. IL-33: A central cytokine in helminth infections. Semin Immunol 53:101532.

9. Cohen ES, Scott IC, Majithiya JB, Rapley L, Kemp BP, England E, Rees DG, Overed-Sayer CL, Woods J, Bond NJ, Veyssier CS, Embrey KJ, Sims DA, Snaith MR, Vousden KA, Strain MD, Chan DT, Carmen S, Huntington CE, Flavell L, Xu J, Popovic B, Brightling CE, Vaughan TJ, Butler R, Lowe DC, Higazi DR, Corkill DJ, May RD, Sleeman MA, Mustelin T. 2015. Oxidation of the alarmin IL-33 regulates ST2-dependent inflammation. Nat Commun 6:8327.

10. Moffatt MF, Gut IG, Demenais F, Strachan DP, Bouzigon E, Heath S, von Mutius E, Farrall M, Lathrop M, Cookson W, Consortium G. 2010. A large-scale, consortium-based genomewide association study of asthma. N Engl J Med 363:1211–1221.

11. Wechsler ME, Ruddy MK, Pavord ID, Israel E, Rabe KF, Ford LB, Maspero JF, Abdulai RM, Hu CC, Martincova R, Jessel A, Nivens MC, Amin N, Weinreich DM, Yancopoulos GD, Goulaouic H. 2021. Efficacy and Safety of Itepekimab in Patients with Moderate-to-Severe Asthma. N Engl J Med 385:1656–1668.

12. Kelsen SG, Agache IO, Soong W, Israel E, Chupp GL, Cheung DS, Theess W, Yang X, Staton TL, Choy DF, Fong A, Dash A, Dolton M, Pappu R, Brightling CE. 2021. Astegolimab (anti-ST2) efficacy and safety in adults with severe asthma: A randomized clinical trial. J Allergy Clin Immunol 148:790–798.

13. Coakley G, McCaskill JL, Borger JG, Simbari F, Robertson E, Millar M, Harcus Y, McSorley HJ, Maizels RM, Buck AH. 2017. Extracellular Vesicles from a Helminth Parasite Suppress Macrophage Activation and Constitute an Effective Vaccine for Protective Immunity. Cell Rep 19:1545–1557.

14. Meiners J, Reitz M, Rudiger N, Turner JE, Heepmann L, Rudolf L, Hartmann W, McSorley HJ, Breloer M. 2020. IL-33 facilitates rapid expulsion of the parasitic nematode. Strongyloides raN from the intestine via ILC2- and IL-9-driven mast cell activation. PLoS Pathog 16:e1009121.

15. Jamwal A, Colomb F, McSorley HJ, Higgins MK. 2023. Structural basis for IL-33 recognition and its antagonism by the helminth effector protein HpARI. bioRxiv doi:10.1101/2023.08.10.552813:2023.08.10.552813.

16. Johnston CJ, Robertson E, Harcus Y, Grainger JR, Coakley G, Smyth DJ, McSorley HJ, Maizels R. 2015. Cultivation of Heligmosomoides polygyrus: an immunomodulatory nematode parasite and its secreted products. J Vis Exp doi:10.3791/52412:e52412.

17. Doellinger J, Blumenscheit C, Schneider A, Lasch P. 2020. Isolation Window Optimization of Data-Independent Acquisition Using Predicted Libraries for Deep and Accurate Proteome Profiling. Anal Chem 92:12185–12192.

18. Neill DR, Wong SH, Bellosi A, Flynn RJ, Daly M, Langford TK, Bucks C, Kane CM, Fallon PG, Pannell R, Jolin HE, McKenzie AN. 2010. Nuocytes represent a new innate effector leukocyte that mediates type-2 immunity. Nature 464:1367–70.

19. Howe KL, Bolt BJ, Shafie M, Kersey P, Berriman M. 2017. WormBase ParaSite - a comprehensive resource for helminth genomics. Mol Biochem Parasitol 215:2–10.

20. Pollo SMJ, Leon-Coria A, Liu H, Cruces-Gonzalez D, Finney CAM, Wasmuth JD. 2023. Transcriptional patterns of sexual dimorphism and in host developmental programs in the model parasitic nematode Heligmosomoides bakeri. Parasit Vectors 16:171.

21. Kouzaki H, Iijima K, Kobayashi T, O’Grady SM, Kita H. 2011. The danger signal, extracellular ATP, is a sensor for an airborne allergen and triggers IL-33 release and innate Th2-type responses. J Immunol 186:4375–87.

22. Bartemes KR, Iijima K, Kobayashi T, Kephart GM, McKenzie AN, Kita H. 2012. IL-33-responsive lineage-CD25+ CD44(hi) lymphoid cells mediate innate type 2 immunity and allergic inflammation in the lungs. J Immunol 188:1503–13.

23. McSorley HJ, Blair NF, Smith KA, McKenzie AN, Maizels RM. 2014. Blockade of IL-33 release and suppression of type 2 innate lymphoid cell responses by helminth secreted products in airway allergy. Mucosal Immunol 7:1068–78.

24. Chauche C, Vacca F, Chia SL, Richards J, Gregory WF, Ogunkanbi A, Wear M, McSorley HJ. 2020. A Truncated Form of HpARI Stabilizes IL-33, Amplifying Responses to the Cytokine. Front Immunol 11:1363.

25. Schiering C, Krausgruber T, Chomka A, Frohlich A, Adelmann K, Wohlfert EA, Pott J, Griseri T, Bollrath J, Hegazy AN, Harrison OJ, Owens BMJ, Lohning M, Belkaid Y, Fallon PG, Powrie F. 2014. The alarmin IL-33 promotes regulatory T-cell function in the intestine. Nature 513:564–568.

26. Hung LY, Tanaka Y, Herbine K, Pastore C, Singh B, Ferguson A, Vora N, Douglas B, Zullo K, Behrens EM, Li Hui Tan T, Kohanski MA, Bryce P, Lin C, Kambayashi T, Reed DR, Brown BL, Cohen NA, Herbert DR. 2020. Cellular context of IL-33 expression dictates impact on anti-helminth immunity. Sci Immunol 5.

27. Bourgeois E, Van LP, Samson M, Diem S, Barra A, Roga S, Gombert JM, Schneider E, Dy M, Gourdy P, Girard JP, Herbelin A. 2009. The pro-Th2 cytokine IL-33 directly interacts with invariant NKT and NK cells to induce IFN-gamma production. Eur J Immunol 39:1046–55.

28. Komai-Koma M, Wang E, Kurowska-Stolarska M, Li D, McSharry C, Xu D. 2016. Interleukin-33 promoting Th1 lymphocyte differentiation dependents on IL-12. Immunobiology 221:412–7.

29. Bonilla WV, Frohlich A, Senn K, Kallert S, Fernandez M, Johnson S, Kreutzfeldt M, Hegazy AN, Schrick C, Fallon PG, Klemenz R, Nakae S, Adler H, Merkler D, Lohning M, Pinschewer DD. 2012. The alarmin interleukin-33 drives protective antiviral CD8(+) T cell responses. Science 335:984–9.

30. Filbey KJ, Grainger JR, Smith KA, Boon L, van Rooijen N, Harcus Y, Jenkins S, Hewitson JP, Maizels RM. 2014. Innate and adaptive type 2 immune cell responses in genetically controlled resistance to intestinal helminth infection. Immunol Cell Biol 92:436–48.

31. Affinass N, Zhang H, Lohning M, Hartmann S, Rausch S. 2018. Manipulation of the balance between Th2 and Th2/1 hybrid cells affects parasite nematode fitness in mice. Eur J Immunol 48:1958–1964.

32. Smith KA, Filbey KJ, Reynolds LA, Hewitson JP, Harcus Y, Boon L, Sparwasser T, Hammerling G, Maizels RM. 2016. Low-level regulatory T-cell activity is essential for functional type-2 effector immunity to expel gastrointestinal helminths. Mucosal Immunol 9:428–43

